# Forecasting hotspots of grassland suitability under climate change for restoration

**DOI:** 10.1101/2024.08.08.607270

**Authors:** Santosh Kumar Rana, Jessica Lindstrom, Melissa A. Lehrer, Marissa Ahlering, Jill Hamilton

## Abstract

● Local species-climate relationships are often considered in restoration management. However, as climate change disrupts species-climate relationships, identifying factors that influence habitat suitability now and into the future for individual species, functional groups, and communities will be increasingly important for restoration. This involves identifying hotspots of community suitability to target seed sourcing and restoration efforts.
● Using ensemble species distribution modeling (eSDM), we analyzed 26 grassland species commonly used in restoration to identify bioclimatic variables influencing their distributions. We predicted habitat suitability under current and future (2050) climates and identified hotspots where diverse species and functional group suitability was greatest. These hotspots of habitat suitability were then overlaid with estimates of landscape connectivity and protected status to quantify potential suitability for restoration now and into the future.
● Temperature and precipitation during warmer quarters largely influenced grassland species habitat suitability. Hotspots of grassland habitat suitability were identified in Minnesota, North Dakota, and South Dakota, with projected northward shifts under future climate scenarios. Overlaying these hotspots with estimates of landscape connectivity and protected status revealed limited connectivity and protection, highlighting regions to prioritize for restoration and conservation efforts.
● Leveraging an understanding of species relationship with climate, this research emphasizes the importance of quantifying connectivity and protected status across aggregated hotspots of species suitability for conservation and restoration. Identifying these hotspots now and into the future can be used to prioritize regions for seed sourcing and restoration, ensuring long-term maintenance of functional ecosystems across grassland communities.

## 1. Introduction

As climate change and land-use modification impact the quality and availability of habitats for species persistence, identifying the factors that influence the success of species or functional groups across diverse and changing landscapes will be critical for restoration (Lyon et al., 2019; Palmer et al., 2016; Tavernia et al., 2013). Restoration often relies upon an understanding of the contemporary climatic conditions influencing species’ presence on the landscape, the climatic niche (Doherty et al., 2017; Potter and Hargrove, 2012; Wilsey, 2021). However, given the pace of global change, shifting climatic conditions may lead to a mismatch in projected species-environment relationships impacting restoration success (IPBES, 2019). Thus, there is a need to identify factors that shape the climatic niche for species or functional groups and to quantify how climate change may impact contemporary species-climate relationships. Leveraging an understanding of the overlap in contemporary and projected future species-climate relationships may aid in identifying habitats to prioritize now and into the future for restoration or conservation.

Ensemble species distribution modeling (eSDM) provides a valuable tool for guiding climate-informed restoration strategies (Zurell et al., 2022). Building upon traditional species distribution models, which quantify species-environment relationships to predict species distributions, eSDMs leverage an ensemble of sub-models to refine predictions of species-environment relationships across diverse spatial resolutions. This approach generates reliable predictions and minimizes uncertainty stemming from variation in individual model predictions, improving the predictive accuracy of fitted models around species-environment relationships (Araújo and New, 2007; Breiner et al., 2015; Grenouillet et al., 2011; Guisan and Thuiller, 2005; Marmion et al., 2009). These models are widely used in conservation and restoration efforts of native species (Butterfield et al., 2016; Eyre et al., 2022; Guisan et al., 2013; López-Tirado and Hidalgo, 2016), determination of seed transfer zones (Bower et al., 2014; Campbell et al., 1991; Doherty et al., 2017), selection of seed sources for restoration (Harrison et al., 2017; Havens et al., 2015; Prober et al., 2015), protection of native species (Shao et al., 2022), and assessing invasion risk (Peeler and Smithwick, 2018; Thuiller et al., 2005). Ultimately, eSDMs offer an integrated approach for forecasting species-environment relationships across different spatial and temporal scales (Sofaer et al., 2019).

Globally, native grasslands are one of the most critically imperiled ecosystems, with much of their historical extent converted to row-crop agriculture (Lindstrom et al., 2023; Volk et al., 2022). This is particularly pronounced in the Northern Great Plains of North America, where up to 87% of historical grasslands have been lost due to agricultural conversion (Comer et al. 2018; Hoekstra et al. 2005; Samson et al. 2004). To mitigate these losses, land managers are actively implementing restoration with a goal to reseed plant communities that will establish, exhibit resilience to change, and persist over time (Lindstrom et al., 2023; Buisson et al., 2022). To target restoration efforts across grassland communities, eSDMs may be used to quantify species-environment relationships and identify regions of projected suitability for diverse community members. Identification of aggregated areas of habitat suitability, defined as hotspots, under contemporary and future climates may provide an effective means to guide restoration and conservation efforts for grassland communities (Zellmer et al., 2019).

Focusing on grassland communities within North America, we leverage an understanding of the climatic niches for 26 grassland species and their functional groups to identify hotspots of habitat suitability based on contemporary and future species-climate relationships. Our study asks (1) What bioclimatic factors influence the contemporary probability of presence and the projected distributions for grassland species commonly used in restoration under global change? (2) and based on current and future climate projections, we ask if there are geographic regions that exhibit substantial overlap in predicted habitat suitability that could be considered aggregated hotspots of climatic suitability for grassland community members? and finally (3) we assess if hotspots of climatically suitable habitat overlap with regions that exhibit connectivity or are protected to support successful restoration and conservation efforts. Ultimately, this research identifies areas of predicted habitat suitability that are or could be targets for restoration or conservation within these anthropogenically-altered landscapes.

## 2. Material and Methods

### 2.1. Focal species and the rarefaction of occurrence points

In this study, we analyzed a total of 26 species that are commonly used in grassland restoration throughout the Northern Great Plains of North America (Table 1; Dixon et al., 2014). These species belong to different functional groups, including 17 forbs, six grasses, and three legumes. Forbs provide several services within grassland ecosystems, including nutritious forage for pollinators and soil stabilization to limit erosion (Siebert and Terblanche, 2020; Siebert and Dreber, 2019). Legumes have the capacity to fix atmospheric nitrogen and facilitate interspecific nitrogen transfer in the ecosystem (Carlsson and Huss-Danell, 2003; Nyfeler et al., 2011). Legumes are also valuable as forage for livestock and can be used as cover crops to reduce erosion and improve soil health. Grasses contribute to carbon sequestration, soil erosion control, and grazing for livestock. Grasses can use available resources, including nitrogen supplied from legumes, to produce high-quality forage (Nyfeler et al., 2011).

**Table 1.**
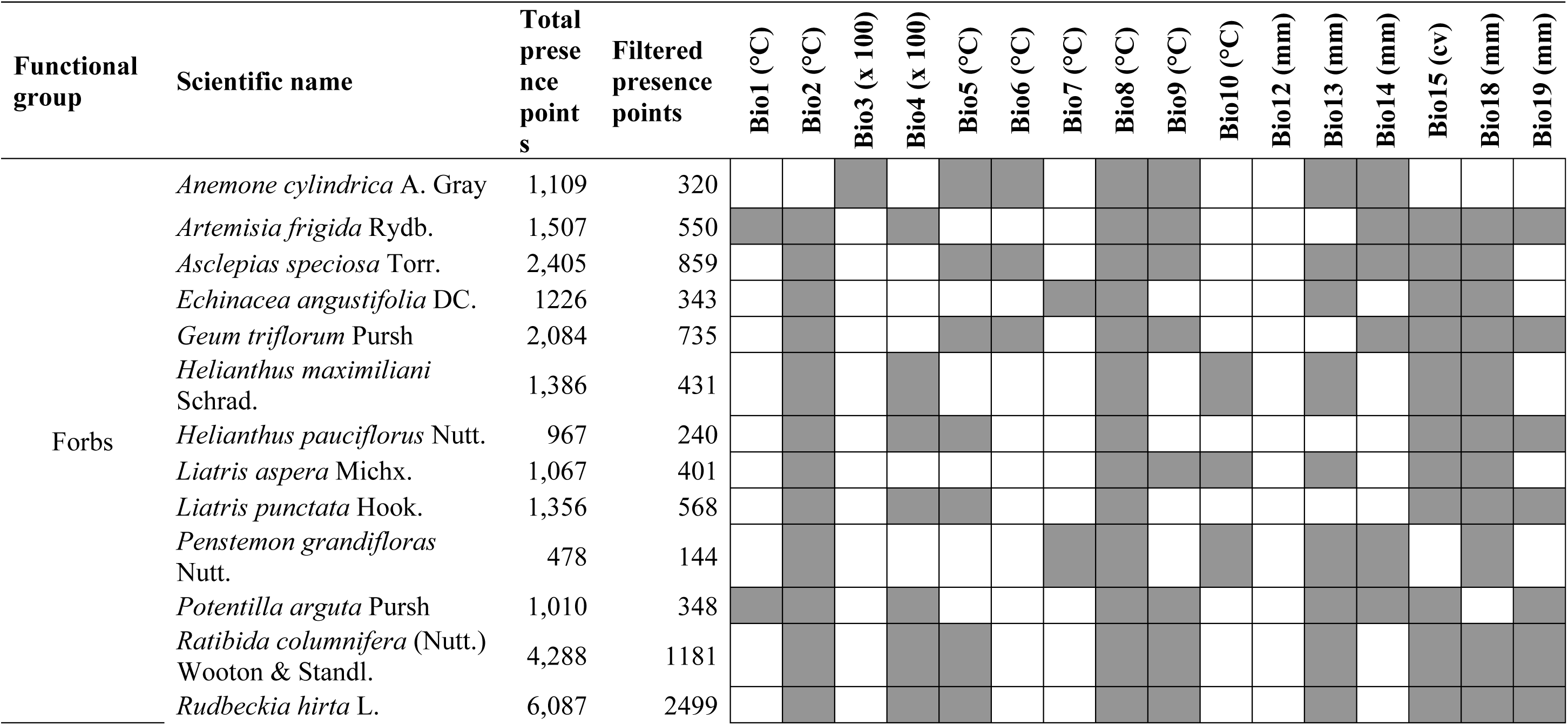
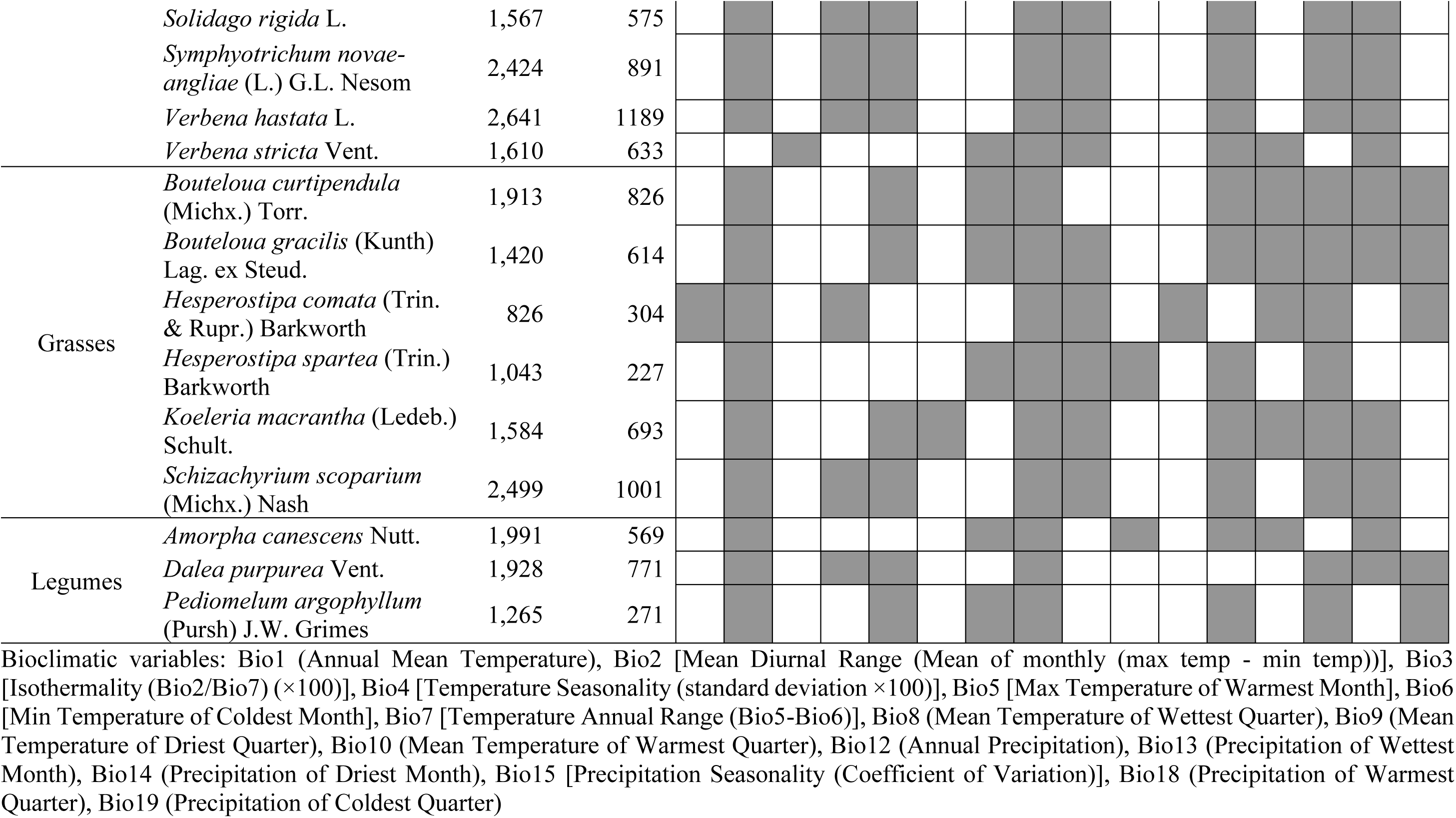
The selected bioclimatic variables (grey highlighted box) for 26 grassland species used in ensemble species distribution modeling (eSDM) using current (1970–2000) and future (2050: 2041–2060) climatic scenarios. eSDM was based on a multi-model median ensemble of general circulation models (24 GCMs; https://www.worldclim.org/data/cmip6/cmip6_clim30s.html) using filtered presence points (out of the total number of presence points) available from ground-validated and Global Biodiversity Information Facility database (GBIF; 05 September 2020; www.gbif.org). Species are grouped by plant functional groups: Forbs, Grasses, and Legumes.

Ground-validated presence locations for all 26 species were first mapped across remnant grassland habitats throughout Minnesota, North Dakota, and South Dakota, USA. Remnant grasslands reflect regions where the soil has not been disturbed due to agricultural activities. Ground-validated occurrence records were supplemented with data extracted from the Global Biodiversity Information Facility database to create occurrence datasets needed for ensemble modeling (GBIF; 05 September 2020; www.gbif.org; Table **1**). GBIF compiles citizen science observations and herbarium records with available geo-coordinates to create contemporary global records for species occurrence. Observations recorded prior to 1900 were excluded to limit the inclusion of false or imprecise location data.

Occurrence datasets associated with GBIF and ground-validated presence locations were used to model the climatic niche of each of the 26 individual species and their functional groups. To minimize the potential impact of overfitting ensemble models, we spatially rarefied geographic occurrences (Boria et al., 2014; Hijmans, 2012; Rana et al., 2021) in a 10 km spatial grid (Boria et al., 2014; Veloz, 2009) using the *spThin* R-package (Aiello-Lammens et al., 2015). The remaining observations were visually filtered, and outliers were removed. In addition, GBIF records that intersected ground-validated locations were omitted. The number of filtered presence points for grassland species ranged from 144 (out of 487 total presence points, *Penstemon grandifloras* Nutt.) to 2499 (out of 6087 total presence points, *Rudbeckia hirta* L.) (Table **1**). In species distribution modeling, pseudo-absence points have been widely used to characterize environments without true absence data (Barbet-Massin et al., 2012). We randomly selected 10000 pseudo-absence points outside a 50 km buffer from the occurrence points. This approach balances presence data with absence data, improving the accuracy of the predicted species distributions.

### 2.2. Predictive variables for ensemble species distribution modeling

The potential contemporary distribution for each of the 26 focal species and three associated functional groups were modeled based on current bioclimatic variables (c. 1970–2000). Bioclimatic variables associated with occurrence points were downloaded from Worldclim (http://www.worldclim.org/version2) at 30-arc sec (∼1 km^2^) resolutions (Fick and Hijmans, 2017). These variables reflect monthly minimum, average, and maximum temperature and precipitation values for 1970 to 2000 associated with each species’ geo-location (Hijmans et al., 2005) (Table 1). Bioclimatic variables are valuable for predicting contemporary species distributions and shifts in response to climate change (Beaumont et al., 2005; Pearson and Dawson, 2003).

To limit the inclusion of correlated or redundant variables, we examined multicollinearity across bioclimatic variables using variance inflation factor (VIF; Fox and Weisberg, 2011). VIF measures the degree to which variance is inflated for the *j*^th^ predictor and is expressed as VIF*j* = 1/(1–R*j*^2^), where R*j*^2^ is the regression coefficient for variable *j* when it is regressed against other predictors, one at a time (Jackson et al., 2009). High multicollinearity inflates the standard errors associated with beta weight which can compromise the reliability and interpretability of regression coefficients. Following the iterative calculation of VIF, which works to eliminate highly correlated variables in a stepwise manner, a group of bioclimatic variables with a VIF less than ten were retained for each species (Quinn and Keough, 2002) (Table S1). Bioclimatic variables were assessed for their contribution to predicting species’ distribution based on their relative importance, derived from the *Biomod2* modeling. The relative importance was scaled to 100%, and bioclimatic variables were reordered for each individual species.

Potential future (2050: average for 2041–2060) distributions were modeled based on current distributions using the ensembled global circulation model (GCM) under a shared socioeconomic pathway (SSP). SSPs represent socioeconomic change derived from projected representative concentration pathways (RCPs), mitigation efforts, and adaptation strategies. We used the middle-of-the-road scenario, SSP2-4.5, one of four scenarios (SSP1-2.6, SSP2-4.5, SSP4-6.0, and SSP5-8.5), to project the current distribution of each grassland species and their functional groups into the future. This scenario uses a moderate radiative forcing level of 4.5 Wm^-2^, which is equivalent to the level used in the IPCC fifth assessment report (AR5) and poses a moderate challenge for both mitigation and adaptation (Fricko et al., 2017; Gidden et al., 2019; Riahi et al., 2017). Future bioclimatic datasets were simulated from WorldClim using 24 downscaled GCMs under SSPs from the coupled model intercomparison project phase 6 (CMIP6). Simulated future bioclimatic datasets were generated using the multi-model median (MMM) ensemble approach for each bioclimatic variable, taking the median values of the available 24 GCMs (Fick and Hijmans, 2017; Herrando-Moraira et al., 2022; Rana et al., 2021). The MMM approach outperforms individual models and accounts for the heterogeneity among several GCMs at global and regional scales (Aguirre-Gutiérrez et al., 2017; Murphy et al., 2004).

### 2.3. Ensemble model development and associated validation in *Biomod2*

Ensemble SDMs were implemented in the *Biomod2* R-package (Thuiller et al., 2020) for each individual grassland species and associated functional groups. *Biomod2* incorporates simulations across multiple sub-models, parameters, and climatic conditions (Araujo and New, 2007). SDMs were assessed for their accuracy and projected using multiple sub-models based on the following modeling approaches: Surface Range Envelop (SRE), Flexible Discriminant Analysis (FDA), General Additive Model (GAM), General Linear Model (GLM), Multivariate Adaptive Regression Splines (MARS), Classification Tree Analysis (CTA), Artificial Neural Network (ANN), Generalized Boosting Model (GBM), Maximum Entropy (MaxEnt), Maximum Entropy new implementation (MaxNet), and Random Forest (RF) (Thuiller et al., 2020). Sub-models were assessed using default parameters within the *Biomod2* package, except for MaxEnt, for which ‘Maximum iteration’ was set as 5000.

Ensemble modeling and validation were carried out in *Biomod2* to identify bioclimatic variables that explain each species’ niche, establishing a consensus for predicted habitat suitability for individual species and functional group based on climatic data. Consensus mapping included an initial calibration that defines a 4-fold cross-validation of the model to partition data into 75% training and 25% evaluation sets. The calibration of the model was performed using 11 sub-models in the ‘*BIOMOD_Modelling*’ function for ten repetitions. The evaluation datasets quantify the predictive power of the model, comparing True Skill Statistics (TSS) and Area Under Curve-Receiver Operating Characteristics (AUC) statistics (Allouche et al., 2006; Araujo et al., 2005). TSS is a threshold-dependent performance measure used to evaluate the accuracy of distribution models. TSS is calculated internally within *Biomod2,* resulting in a matrix that assesses the probability that presence and/or absence points are accurately predicted in eSDM (Allouche et al., 2006). TSS values range from −1 to +1, where +1 indicates a perfect prediction, 0 indicates no improvement over random prediction, and -1 indicates total disagreement between observed and predicted values (Allouche et al., 2006). In addition, we used AUC, a threshold-independent performance measure, to evaluate model performance; AUC is particularly suitable for evaluating the performance of ordinal score models such as logistic regression (Phillips et al., 2006). A model’s predictive accuracy is deemed average when the AUC value is below 0.7, high when it falls between 0.7 and 0.9, and excellent when it is greater than or equal to 0.9 (Phillips et al., 2006; West et al., 2016).

We applied the ‘*BIOMOD_EnsembleModeling*’ function for those sub-models with TSS > 0.7, except for two of the species, *Koeleria macrantha* (Ledeb.) Schult. and *Schizachyrium scoparium* (Michx.) Nash, for which we applied TSS > 0.65. When calibrating the initial model for these two species, only one or two sub-models achieved a TSS value greater than 0.7, making it impossible to include multiple sub-models for ensemble modeling. Thus, a threshold was set for these two species, allowing multiple sub-models to be used in ensemble species distribution modeling. Using ‘*BIOMOD_Projection*’, we projected calibrated species distributions using contemporary bioclimatic variables into geographic space. Lastly, the ‘*BIOMOD_EnsembleForecasting*’ function was used to forecast and generate the predicted habitat suitability consensus for each individual grassland species and associated functional groups.

### 2.4. Ensemble species distribution modeling for grassland species and associated functional groups

The ensemble model was converted to a binary presence/absence (i.e., suitable/unsuitable) map based on individual species and functional group distributions. This conversion used a 30% threshold of the projected habitat suitability to avoid over-projection (Forester et al., 2013; Rana et al., 2020, 2021) and was used to fit the contemporary distribution of individual grassland species and their functional groups. Pixel counts of the raster layers were converted into suitable habitat and expressed as km^2^ and percentage of suitable habitat out of the total modeled extent of North America (i.e., 24.7 million km²). Using ArcMap 10.4.1 (ESRI, 2016), the predicted spatial extent of grassland species and functional groups was categorized into reduced, stable, or expanding based on a comparison between current and future projections (2050). ‘Reduction’ represents a decrease, ‘Stable’ represents no change, and ‘Expansion’ represents an increase in the predicted area of suitable habitat under global change.

### 2.5. Aggregating contemporary and future habitat suitability to identify hotspots of suitable grassland habitat and their connected and protected status

Hotspot analysis was employed to identify geographic regions where predicted habitat suitability overlapped for multiple species within a grassland community under both current and future climatic conditions (O’Donnell et al., 2012). Following O’Donnell et al. (2012), we aggregated the individual binary maps of the climatically suitable niches for all 26 species using the Raster Calculator within the Spatial Analyst Extension of ArcGIS 10.4.1. The resulting aggregated map revealed the geographic extent of overlapping habitat suitability for various combinations of species. Areas where at least 25% of the aggregated map of suitability was predicted to support at least eight out of the 26 studied grassland species were defined as hotspots (O’Donnell et al., 2012; Shrestha et al., 2022). These hotspots were further categorized based on species richness into three categories: 1) 8 to 13 species, 2) 14 to 19 species, and 3) 20 to 26 species.

To quantify and visualize landscape connectivity for predicted hotspots of habitat suitability, we evaluated the overlap between hotspots under current and future scenarios with the assessment of connectivity from the Resilient & Connected Networks (RCN) map created by The Nature Conservancy (https://maps.tnc.org/resilientland/; Anderson et al., 2023). The RCN map integrates three nationwide assessments of habitat resilience, landscape connectivity, and biodiversity significance that may be used to target preservation or restoration efforts. We used the landscape connectivity category to assesses landscape features associated with connectivity and climate flow. Under the expectation that geophysical features remain stable over time, we used the current RCN map to extract regions overlapping with hotspots of habitat suitability for both current and future climate scenarios. These shared regions were extracted using the ‘Zonal Statistics as Table’ function of spatial analyst in ArcGIS.

To evaluate the protected status of the hotspots, we extracted the overlap between hotspots of habitat suitability with a data layer describing the protected status for North America, as designated by the International Union for Conservation of Nature (IUCN) (CEC, 2021). Extracting this overlap is crucial for identifying gaps in current conservation efforts. Overlap between hotspots of habitat suitability with landscape connectivity and IUCN’s protected area categories were calculated as a percentage (%) of the total modeled area (total area of North America, 24.71 million km²).

## 3. Results

### 3.1. Bioclimatic factors contributing to species and functional group climatic niche

Following assessment of bioclimatic variables used to predict species and functional group distributions, we identified a set of uncorrelated variables based on the VIF (VIF*j*<10) (Tables 1 and S1) that contribute to predictions of habitat suitability. Of these variables, ‘Max Temperature of Warmest Month (bio5)’ and ‘Precipitation of Warmest Quarter (bio18)’ contributed significantly to predictions for 16 and 21 grassland species, respectively (Fig. 1).

**Figure 1.**
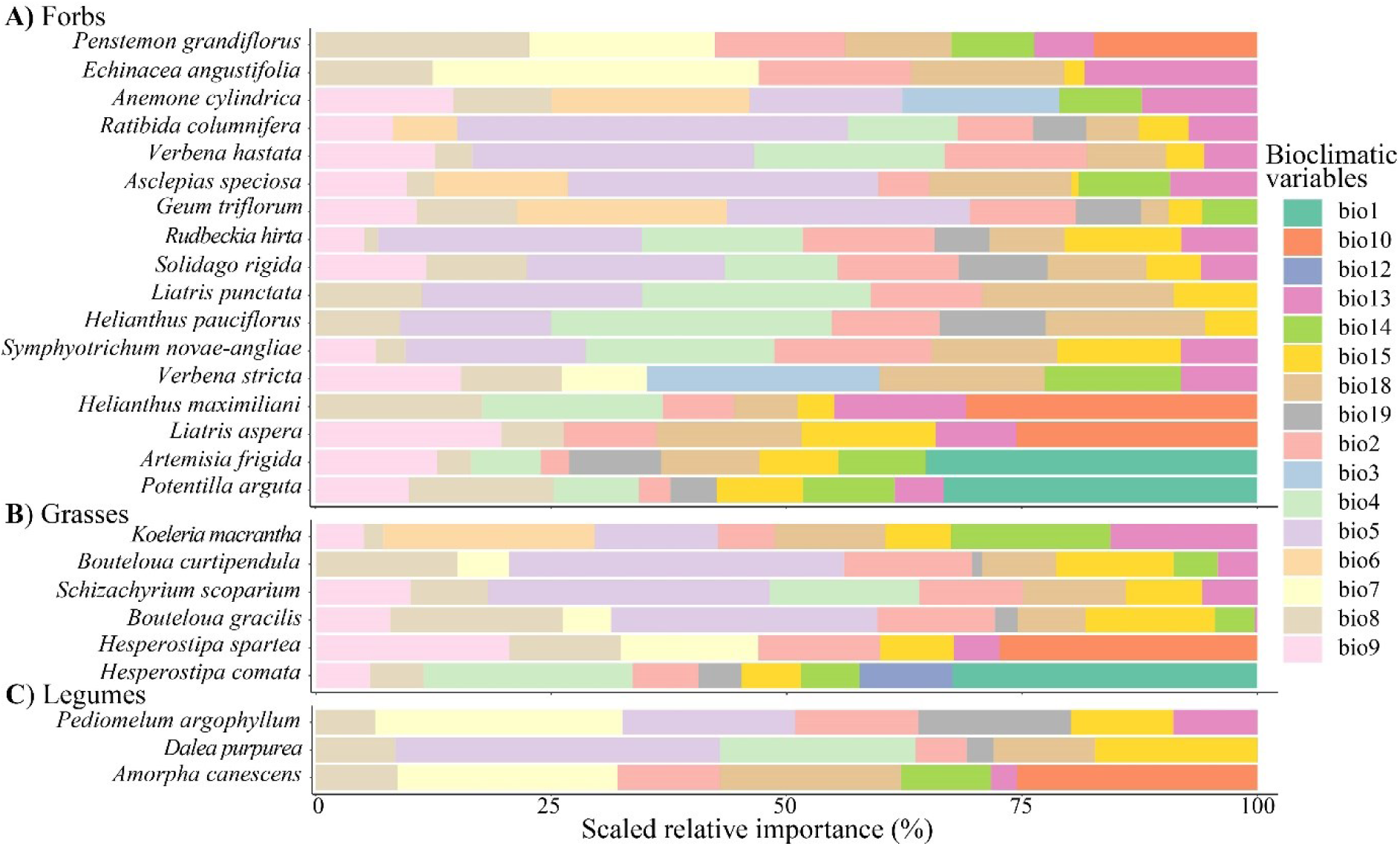
The scaled relative importance (%) of bioclimatic variables for the suitability of 26 functionally categorized grassland species (**A**, Forbs; **B**, Grasses; and **C**, Legumes) used in ensemble species distribution modeling (eSDM). The percentage of each variable’s importance is scaled to 100% and reordered according to the relative importance within each functional group. Refer to Table S1 for the abbreviated bioclimatic variables.

For most forbs, at least nine distinct bioclimatic factors were required to forecast predicted distributions (Fig. 1 and Table 1). This indicates that a diverse range of bioclimatic factors are needed to accurately forecast habitat suitability. However, select species like *Liatris punctata* Hook., could be effectively predicted using six bioclimatic factors (Fig. 1). On average, the projected distributions of the forb species appeared to be largely influenced by temperature-determinants over precipitation variables. The predicted distributions for most species within the grasses functional group were determined by temperature-related variables, with ‘Max Temperature of Warmest Month’ (bio5) and ‘Mean Temperature of Driest Quarter’ (bio9) key factors contributing to habitat suitability predictions within that functional group (Fig. 1 and Table 1). For legumes, at least six different bioclimatic variables were required to predict species distributions, including ‘Mean Diurnal Range’ (bio2) and ‘Mean Temperature of Wettest Quarter’ (bio8), which contributed most significantly to the modeled distribution of legumes (Fig. 1). Interestingly, when comparing across functional groups ‘Mean Temperature of Driest Quarter’ (bio9) appeared to contribute significantly to the projected distribution of forbs and grasses, but not legumes.

### 3.2. Predicted shift in suitability for grassland species and associated functional groups under global change

Species- and functional-group specific eSDMs with sub-models of high accuracy (Fig. S1) were used to predict contemporary suitability as well as shifts in the suitable distributions of grassland species under future climates. The comparison in distributions between current and future climates revealed a change to predicted suitability and potential shifts in the predicted distribution of species and functional groups in the future (Figs. 2, 3, S2, and S3).

**Figure 2.**
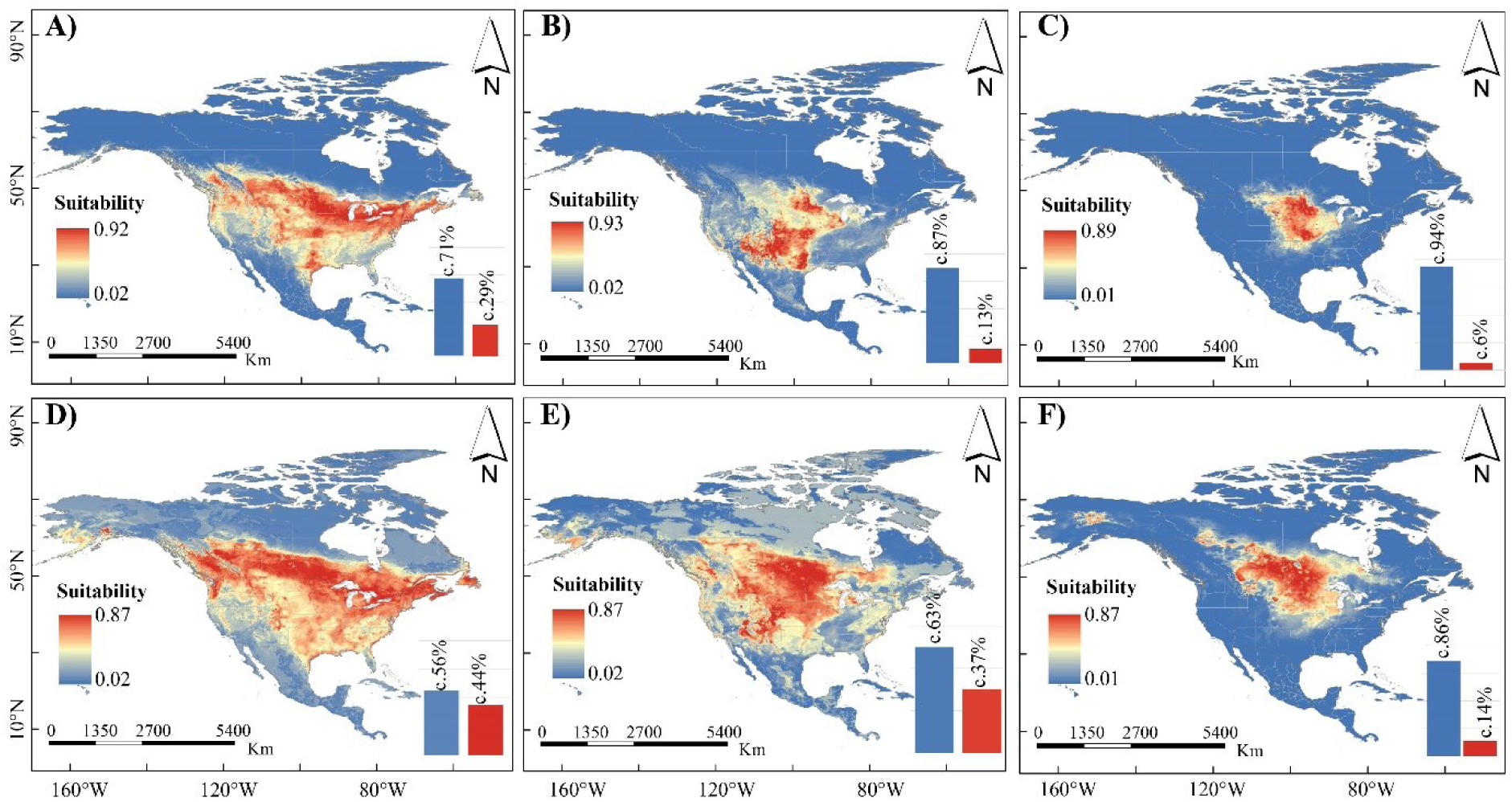
Predicted habitat suitability for the three functional groups (**A, D**) Forbs, (**B, E**) Grasses, and (**C, F**) Legumes under the current (**A**–**C)** and future (**D**–**F**) bioclimatic variables’ scenarios. The legend on the left side of each map shows habitat suitability ranging from high-suitable (red) to low-unsuitable (blue). The percentage of the projected habitats defined as suitable (red bar) and unsuitable (blue bar) are based on a 30% threshold of the projected habitats; these areas are displayed as the bar graph in the lower right corner of each figure. The numbers on top of the bar graph represent the percentage (%) of suitable area out of the total modeled area (i.e., total area of North America, 24.71 million km²).

**Figure 3.**
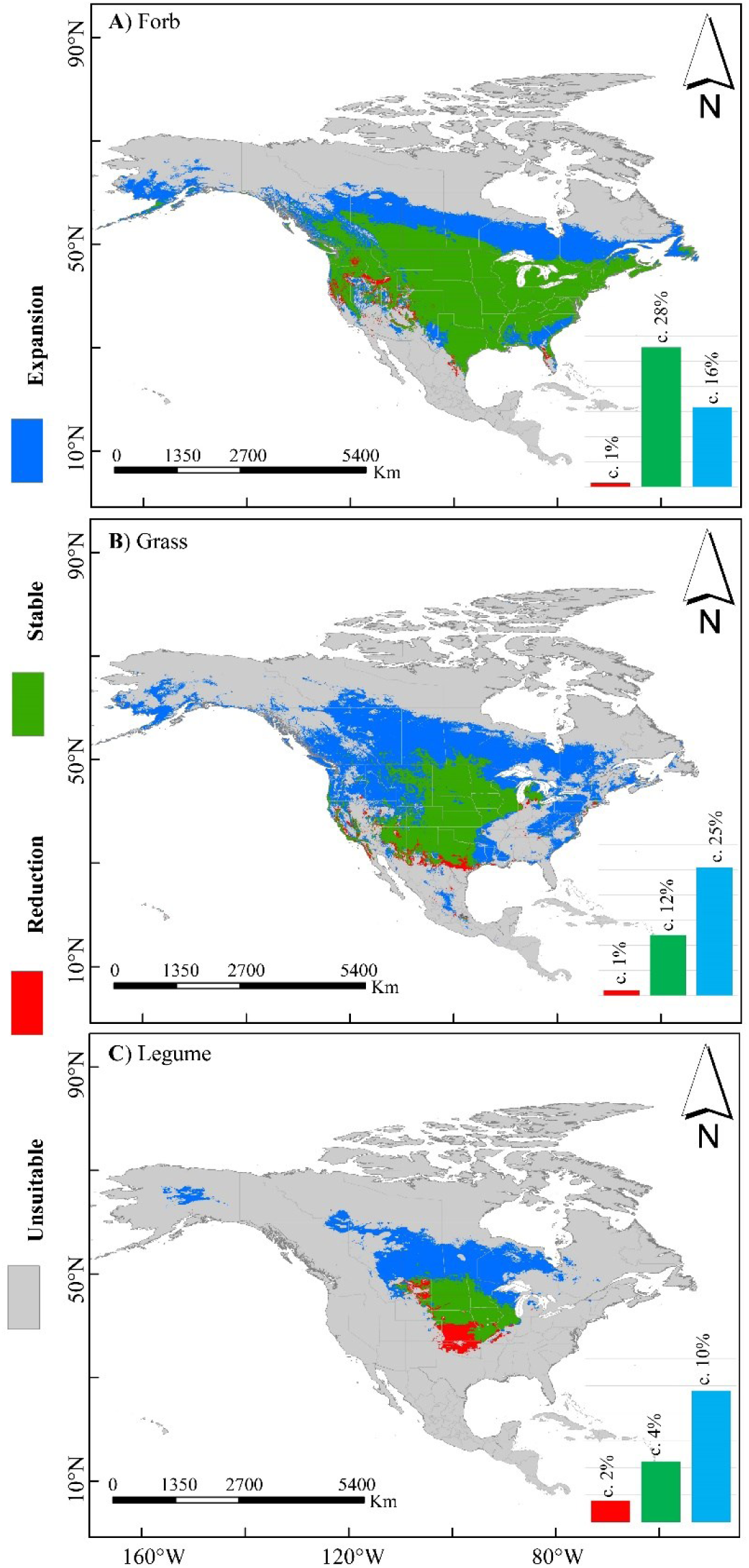
Predicted habitat suitability for functional groups classified as reduced, stable, and expanded areas for (**A**) Forbs, (**B**) Grasses, and (**C**) Legumes. The three categories of suitable habitat calculated as a function of raster aggregation for the current and future bioclimatic variables in the eSDM projections were classified as ‘Reduction’ representing a decrease (red), ‘Stable’ representing no change (green), and ‘Expansion’ representing an increase (blue) in predicted suitable habitat area in the future (2050) as compared to the current scenario. The bar graph in the lower right corner represents the percentage (%) of classified suitable habitat area out of the total modeled area (i.e., total area of North America, 24.71 million km²).

The predicted geographic distribution for forbs was largely concentrated across central and eastern North America, approximately 29% of the modeled area (Figs. 2A, S2 and S4A). However, while most of the projected distributions of forbs were predicted to remain stable (28% of the modeled area) in 2050 (Fig. 3A), there was a projected increase of 16% of the modeled area under future climatic conditions. Thus, despite substantial stability in the projected area of forbs under future conditions, climatically suitable areas are also expected to expand northward in the 2050 scenario (Figs. 2D, 3A, S3 and S5). Among the forbs, *R. hirta* exhibited the largest area of projected suitable habitat, nearly 47 times greater than that of the species with the least suitable area, *P. grandiflorus* (Figs. S2 and S4A). While the predicted habitat suitability for most forb species concentrates in central and eastern North America, *Artemisia frigida* Rydb., *Asclepias speciosa* Torr., and *Geum triflorum* Pursh exhibit geographically dispersed distributions throughout the eastern and western regions of the United States (Fig. S2).

Grass species suitability was predominantly predicted for the central region of North America, with a fragmented distribution of suitability in the eastern region of the United States, occupying 13% of the modeled area based on contemporary climatic conditions (Fig. 2B). Considering 2050 climates, 37% of the modeled area was projected to be suitable, but with increased fragmentation. Indeed, there was a projected shift in suitability, with 25% of the modeled area shifting towards the northwestern United States by the year 2050 (see Figs. 2E, S4, and S5). This shift indicates a likely increase in habitats that could support diverse grass species, especially those sharing similar ecological roles within the community. Of the grass species evaluated, *Bouteloua curtipendula* (Michx.) Torr. exhibited the largest extent of climatically suitable habitat (5% of the modeled area), indicating its broad climatic niche when compared to the other species within the functional group (comprising 2.4–4.8 % of the modeled area) (Figs. S2U and S4A). *Bouteloua curtipendula* is projected to retain the largest area of suitable habitat into the future compared to the other grass species, expanding to encompass 12% of the modeled area with a notable northward shift of approximately 8.59% of the modeled area (Fig. S4B).

In general, the predicted contemporary distribution for legumes remained in the central United States, covering approximately 6% of the modeled area (Figs. 2, S2 and S4A). However, habitat suitability was projected to increase towards central North America under future climates (Fig. 2F), with an approximately 8% increase in suitable habitat (Figs. 2F and S4). Among different legume species, *Dalea purpurea* Vent. Barneby stands out as having the largest area of projected habitat suitability and a region of predicted suitability three times greater than the area of suitability projected for *Pediomelum argophyllum* (Pursh) J.W. Grimes, which has the least projected suitable habitat (Fig. S4A). In addition to having the greatest area of projected habitat suitability under current conditions, *D. purpurea* is predicted to have the greatest area of projected suitable habitat in the future, shifting northward and occupying about 6% of the modeled area in 2050 (Figs. S2, S3, S4B and S5).

### 3.3. Aggregating species-rich hotspots of suitability for resilient and connected landscapes

We identified hotspots of suitability, defined as suitable habitat for diverse grassland community members, and categorized them based on the species suitable for hotspot region. The threshold to identify ‘hotspots of habitat suitability’ were based on ‘25% of the aggregated map of suitability’ requiring aggregation of a minimum of 8 out of 26 species projected distributions (Fig 4A, D). For an area to be considered a hotspot, a minimum of 8 species were required to be suitable for the region, but the composition of these 8 species could vary. Using these parameters, 3% of the modeled area in North America was classified as a ‘hotspot’ of grassland community suitability based on contemporary projections across the 26 species’ climatic niches (Fig. 4D, F). Species-rich hotspots, where the suitable habitat is optimized for the maximum number of grassland species (20 to 26 species), occurred across approximately 0.1% of the modeled area within the tallgrass prairie of the Midwest region of the United States (North Dakota, South Dakota, and Minnesota) (Fig. 4A, C). For the different levels of species richness (i.e. for >8 species), Midwestern states were considered a species-rich region and a hotspot of grassland suitability (Fig 4A).

**Figure 4.**
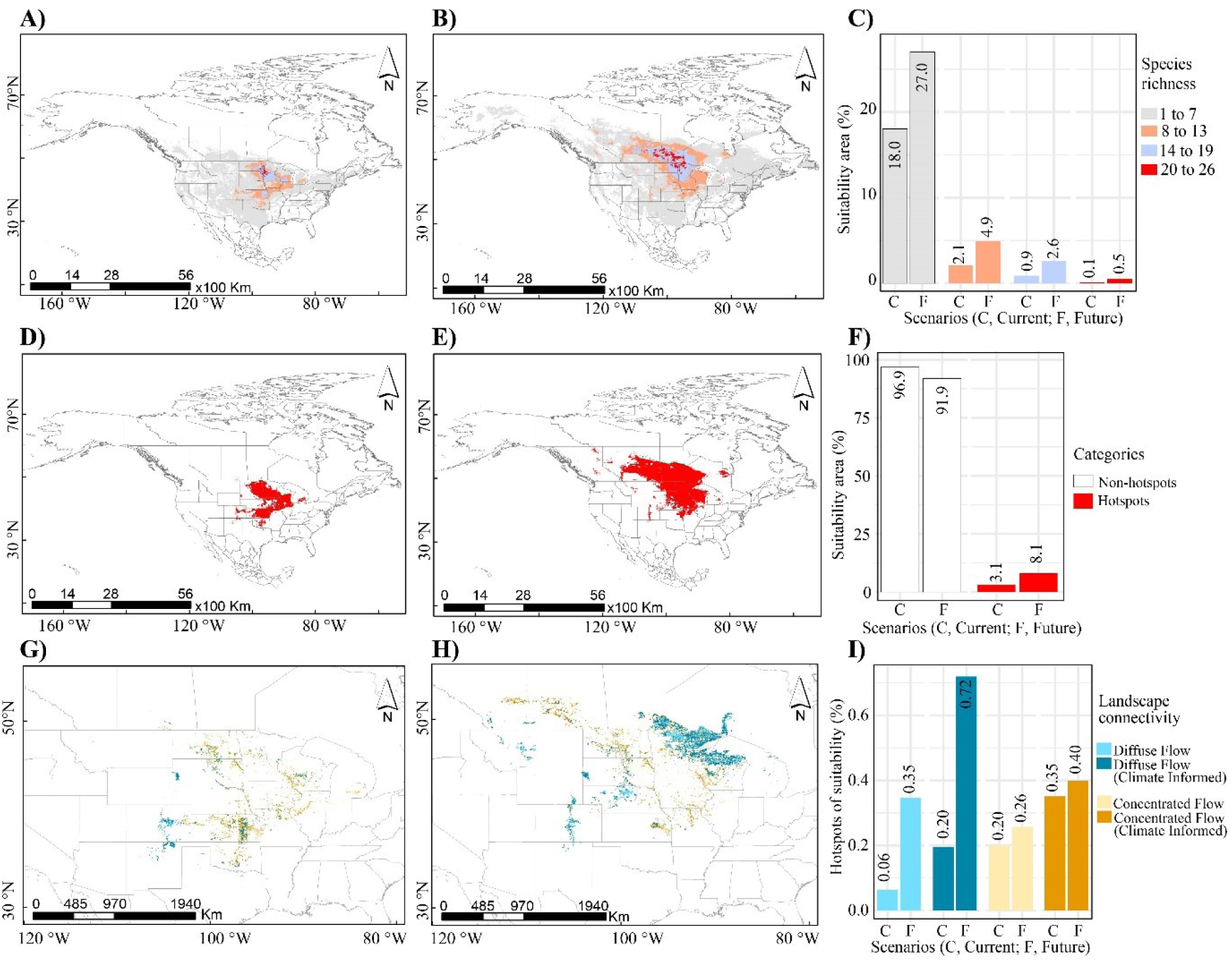
Predicted suitable grassland habitat classified as species richness (**A**–**C**), hotspots of suitable grassland habitat that optimizes the suitable habitat for multiple grassland species (**D– F**), and the overlapped hotspots of suitability with landscape connectivity of resilient and connected networks (RCN) map (**G**–**I**), under the current [C] (**A**, **D**, **G**) and future [F] (**B, E, H**) bioclimatic scenarios. The percentage of predicted suitable habitat areas classified as species richness is expressed in the bar graph highlighting the species composition required to form the grassland suitability (**C**). The percentage of predicted hotspots of suitable habitat areas is expressed in the bar graph (**F**). The resilient and connected network maps are redrawn based on the overlapped hotspots of suitability and landscape connectivity of The Conservancy’s RCN map (The Nature Conservancy, 1975, 2020; https://maps.tnc.org/resilientland/) (**I**). The expressed suitable habitat area (%) is out of the total modeled area (i.e., total area of North America, 24.71 million km²).

Given future climate projections, models indicated that the projected hotspot of suitability for more than eight species will increase to 8% of the modeled area, resulting in a 5% increase from the contemporary projection (Fig. 4E, F). Northern portion of the species’ distributions in Canada exhibited the greatest shift in projected hotspot, with Saskatchewan, Manitoba and Ontario exhibiting projected core hotspots of habitat suitability (for 20 to 26 species) (Fig. 4B). However, projections indicated that North Dakota and Minnesota would likely remain hotspots of suitable grassland habitat as they included projected suitability for the largest number of grassland species (Figs. 4B). Notably, future projections suggest hotspots will become increasingly confined to distinct geographic regions, which may point towards an increased impact of fragmentation long term.

Following identification, hotspots were mapped onto the landscape connectivity map of The Nature Conservancy’s Resilient and Connected Network emphasizing the ‘concentrated flow’ and ‘diffuse flow’. The regions categorized as ‘concentrated flow’ and ‘diffuse (high) flow’ represent narrow and more expansive connectivity pathways that facilitate species movement, respectively. Overlap with the RCN’s connectivity categories suggests that approximately 0.55% of the predicted contemporary hotspots overlap with regions defined as ‘concentrated flow’. In contrast, approximately 0.26% of the projected grassland community hotspot aligns with regions of ‘diffuse flow’ (Fig. 4G–I). This lack of overlap suggests that only a small fraction of the predicted hotspots of grassland community suitability coincide with regions experiencing connectivity or ‘diffuse flow’ exacerbating the impacts of fragmentation in these regions.

We used the existing, contemporary landscape connectivity map of the RCN to extract values that will overlap with hotspots of suitability under future climates. The changes in hotspots of suitability are partly reflected in the RCN’s landscape connectivity categories, especially the anticipated northward movement of the grassland suitability hotspot towards Canada by 2050. Projections indicated a notable increase in the overlap of projected hotspots with (approximately 0.92% of the modeled area) landscape connectivity based on concentrated and diffuse flow compared to contemporary projections (Fig. 4D). This suggests distinct regions of suitability may be more connected under expanding distributions.

Finally, we evaluated the overlap between projected hotspots of grassland suitability and designated protected areas status as defined by IUCN. Our findings suggest there is minimal overlap between the contemporary hotspots of habitat suitability and designated protected areas, with the overlap accounting for less than 0.2% of the modeled area (Table 2). However, this overlap is projected to increase by 0.56% in the future, with ‘National Parks’ projected to experience the highest increase (c. 0.19% of the modeled area) by 2050 (Table 2).

**Table 2.** Predicted hotspots of suitability based on current and future (2050) bioclimatic projection of 26 grassland species overlapped with the protected areas in North America, as designated by the International Union for Conservation of Nature’s (IUCNs) (CEC, 2021). The hotspots of grassland suitability were based on the chosen threshold of 25% that emphasizes regions where at least 25% or more of the aggregated suitable habitat of 26 grassland species is deemed suitable for supporting a diverse array of grassland species. Note: The expressed suitable habitat area (%) is out of the total modeled area (i.e., total area of North America, 24.71 million km²).

| IUCN Classification of the protected area | Hotspots of suitable habitat [km<sup>2</sup> (%)] |  |  |
| --- | --- | --- | --- |
|  | Current scenario | Future scenario | Change |
| Strict Nature Reserve (Ia) | 42.56 (<0.01%) | 2539.33 (0.01%) | 2496.77 (0.01%) |
| Wilderness Area (Ib) | 1205.83 (<0.01%) | 44232.57 (0.18%) | 43026.74 (0.18%) |
| National Park (II) | 56.74 (<0.01%) | 47750.75 (0.19%) | 47694.01 (0.19%) |
| Natural Monument or Feature (III) | 70.93 (<0.01%) | 709.31 (<0.01%) | 638.38 (<0.01%) |
| Habitat/Species Management Area (IV) | 2411.65 (0.01%) | 13902.48 (0.06%) | 11490.83 (0.05%) |
| Protected Landscape/Seascape (V) | 33422.69 (0.14%) | 44601.41 (0.18%) | 11178.72 (0.04%) |
| Protected area with sustainable use of natural resources (VI) | 11419.89 (0.05%) | 34798.75 (0.14%) | 23378.86 (0.09%) |

Note: The expressed suitable habitat area (%) is out of the total modeled area (i.e., total area of North America, 24.71 million km<sup>2</sup>).

## 4. Discussion

In this study, ensemble species distribution modeling was used to assess habitat suitability for grassland species under current and future climatic scenarios, identifying habitats for targeted restoration of individual species, functional groups, and grassland communities. On average, ‘Max Temperature of the Warmest Month (bio5)’ and ‘Precipitation of Warmest Quarter (bio18)’ had the largest influence on the modeled species’ distributions. However, there was substantial individual and functional group variance in the predictors of habitat suitability, indicating the importance of evaluating the climatic factors contributing to individual and functional group habitat suitability. Identifying suitable habitats and shifts in suitable habitats identified hotspots of habitat suitable for diverse grassland community members. In addition, determining how those hotspots intersect with landscape connectivity networks and protected areas identified potential gaps in contemporary preservation and opportunities to enhance restoration to improve connectivity. Ultimately, integrating information on hotspots of projected grassland suitability with connected and conserved networks will inform restoration priorities.

### 4.1. Bioclimatic variables influence the grassland species suitability

Effective restoration relies upon an understanding of the complex relationships between species with climate now and into the future. Using ensemble species distribution modeling we used bioclimatic variables, to predict the current and future species-climate niche and functional group-climate niche to inform targeted restoration. The differential impact of bioclimatic variables to species’ projected distributions highlights their diverse roles in shaping the distributions of species and functional groups across communities and points towards the importance of tailoring habitat suitability assessments to the unique climatic needs of each species or group (Bagne et al., 2012; Havrilla et al., 2023; Jones et al., 2019; Martinson et al., 2011; Volenec and Belovsky, 2018; Zhu et al., 2016).

Many bioclimatic variables were required to predict the distribution of the forb functional group, suggesting the diversity of species within this broad functional group may impact projections. Forbs demonstrate a broad spectrum of adaptations to various environmental conditions, a diversity that has evolved in response to evolutionary pressures like herbivory, pollination strategies, and habitat preferences (Dormann et al., 2001). As a result, to accurately predict their distribution, it is necessary to use a broad set of bioclimatic factors (Austin, 2007; Turner and Knapp, 1996). However, some forbs, like *L. punctata*, require fewer variables, indicating that individual species themselves may exhibit more specialized ecological niches or are adapted to a narrower range of conditions (Martinson et al., 2011).

In contrast, grasses and legumes required fewer bioclimatic variables, indicating less individual variation among these functional groups. Notably, temperature variables were particularly crucial for these functional groups, emphasizing the important role of temperature as a factor influencing life history and function within these groups, including photosynthesis, growth, reproduction, and overall survival (Blair et al., 2014; Brotherton and Joyce, 2015). This suggests that temperature is crucial across diverse grassland species, but detailed focus on targeted species may be necessary for effective conservation and ecosystem management.

### 4.2. Grassland communities are projected to shift northward in response to global change

On average, our model indicates a potential northward expansion of suitable habitat for grassland community members under future climate scenarios (Bagne et al., 2012; Frelich and Reich, 2010; Friggens et al., 2012; Lyon et al., 2019; Martinson et al., 2011). Despite these projections, this does not necessarily imply that grassland species will move northwards (Bagne et al., 2012; Gray and Hamann, 2013; Iverson and Prasad, 2002). Firstly, the ability of species to migrate depends on various factors beyond climatic suitability, such as physical barriers, ecological interactions, and human land use (Knowlton and Graham, 2010; Lyon et al., 2019; Martinson et al., 2011; Tomiolo and Ward, 2018). Many of the regions we modeled have been converted to row-crop agriculture or border the boreal forest ecosystem, which could impede their predicted northward movement (Chapin III and Danell, 2001; Frelich et al., 2024). Secondly, species migration depends on biological factors like dispersal ability and adaptation, which may vary among the diverse species included in this work (Lyon et al., 2019; Tomiolo and Ward, 2018). Therefore, while our predictions suggest a northward shift of suitable grassland habitat, the actual movement of grassland species may not align with these predictions. This discrepancy highlights the need for caution interpreting future models of projected species distributions.

In addition, understanding how different functional groups within grassland communities respond to climate change is important. Contrary to the assumption that similar functional groups would have uniform responses, this study suggests that these groups are projected to respond differently to climate change (Chen et al., 2017; Corlett and Westcott, 2013). Diverse responses across functional groups could stem from unique species characteristics, ecological preferences, habitat linkages, human influences, and evolutionary backgrounds (Aitken et al., 2008; Leimu and Fischer, 2008; Nathan, 2006; Nathan and Muller-Landau, 2000; Schupp et al., 2010; Sullivan et al., 2020). Grasses, for instance, may face significant habitat loss in some areas. This could lead to a decline in grass populations or shifts in their geographical distribution. Forbs, with their greater niche breadth, might either retain their current range or expand. Legumes, however, show a more restrained northward movement, possibly due to their specific habitat requirements or reduced plasticity to respond to rapid climatic shifts (Harrison et al., 2013). In summary, these findings illustrate the varied effects of climate change on different species within grassland ecosystems, pointing to complex implications for ecosystem structure, plant community composition, and conservation strategies.

### 4.3. Grassland hotspots reflects the species-rich area to safeguard grassland

Our study aimed to identify hotspots of habitat suitable for grassland community members, areas with favorable climatic conditions suitable for a diverse range of grassland species (Risser, 1988). Using eSDM, we successfully identified regions that in aggregate are suitable for a rich diversity of grassland species. These relatively small, but ecologically significant hotspots (about 3% of the modeled area) identify habitats suitable for a minimum of 8 grassland species. The Midwest United States, including North Dakota, South Dakota, and Minnesota, emerged as a core hotspot area for a high number of species (ranging from 20 to 26 species), despite covering only 0.1% of the modeled area. This compact biodiversity hotspot is ecologically vital yet vulnerable, underscoring the need for conservation efforts to protect these regions which are projected to support diverse grassland community members. In addition, our projections suggest that hotspots of grassland suitability are shifting northward, driven by moderate CMIP6 warming scenarios (Eyring et al., 2016; Sala et al., 2000), potentially creating new regions favorable for the growth and development of multiple grassland species (Sala et al., 2000). However, even with this predicted northward shift North Dakota and Minnesota are expected to remain core hotspots, suggesting proactive planning for conservation and habitat restoration within these regions will have long-term benefits.

Potential limitations to modeling hotspots of diverse species suitability assumes no interspecific competition (Chase and Leibold, 2003; Wandrag et al., 2023). Competition can significantly impact species distribution and abundance, potentially excluding some species from their projected suitable habitats. Considering biotic interactions like competition and predation alters the realized niche of community members (Hutchinson, 1975; Wiens et al., 2009). Consequently, the study maximizes the identification of the fundamental niche, or habitats that could potentially support multiple grassland species, representing potential hotspots of suitability. In an ideal scenario, the approach outlined here identifies habitats that could support diverse species and may be targets for restoration or preservation to ensure maintenance of functioning grassland ecosystems long term.

### 4.4. Grassland hotspots represents the protected and connected landscapes needed for species movement

This study highlights the importance of connectivity in supporting grassland resilience, and the role connectivity and preservation of potential hotspots of suitable habitat may play in maintaining functional grassland ecosystems. Connectivity ensures genetic exchange and dispersal critical to countering the impacts of fragmentation and demographic loss across populations (Christie and Knowles, 2015; Keeley et al., 2018). Overlaying The Nature Conservancy’s RCN map with hotspots of grassland community suitability we quantified the degree of potential connectivity across regions that should promote diverse communities (Wimberly et al., 2018). We found two types of overlap: ‘concentrated flow’ and ‘diffuse (high) flow’, representing narrow and expansive pathways for species movement, respectively. Overlay analysis with The Nature Conservancy’s RCN map reveals that our identified hotspots of grassland community suitability exhibit limited connectivity. Specifically, only 0.55% of these hotspots show ‘concentrated flow’, and an even smaller 0.26% align with ‘diffuse flow’ regions, suggesting significant fragmentation. This minimal overlap indicates that current connectivity is inadequate, presenting challenges for species dispersal and adaptation in response to climate change. The data highlights a pressing need for strategies to enhance linkage between these hotspots to maintain ecosystem resilience and support diverse communities. Addressing the connectivity gaps is crucial for the long-term stability of grassland ecosystems, especially in the context of evolving environmental conditions.

Projected climate scenarios indicate a tendency for grassland suitability hotspots to migrate toward regions characterized by ‘diffuse flow’. This observed trend underscores a concrete impact of climate change, which is reshaping the habitable landscapes for grassland species, altering how and where these species can thrive within interconnected ecosystems. Our data reflect this shift, revealing that the areas currently facilitating broad species dispersal are likely to become increasingly crucial as climate-induced changes in habitat suitability become more pronounced. Therefore, our landscape connectivity assessments must integrate these climate projections to effectively prioritize and manage conservation efforts for enduring grassland biodiversity. This data can also be used to target regions that may ensure connectivity in the future to minimize the effects of fragmentation to the maintenance of these biodiverse communities.

Finally, the limited overlap of hotspots of species suitability with protected status emphasizes the call for expanded conservation efforts in these regions, particularly to ensure connectivity is maintained across hotspots of biodiversity. While future projections indicate National Parks may comprise a greater percentage of hotspots of suitable grassland habitat, our data also suggests the need to expand or establish new protected regions to ensure maintenance of connectivity across fragmented landscapes. Together these data aid in identifying regions where conservation and restoration efforts will maintain hotspots of grassland suitability critical to restoration that may persist in a connected and preserved landscape.

## 5. Conclusions

The use of ensemble species distribution modeling (eSDM) has provided a valuable perspective on the suitability of habitats for grassland species under diverse climatic conditions, offering a means to strategize regions for conservation and restoration. We have identified hotspots of projected suitable habitat that optimize the habitats suitable for a diversity of species under ideal scenarios. Our findings highlight hotspots of habitat suitable for restoration and preservation, particularly in regions where suitable habitats align with landscape connectivity and protected areas. Our models predict a notable northward shift in the projected distribution of grassland communities due to climate change, signaling the need for adaptive management that considers the potential for dispersal and connectivity in a changing climate. Ultimately, quantifying species- and functional group relationships with climate and identifying hotspots of habitat suitable for diverse grassland community members will allow us to make informed restoration and conservation decisions to ensure the long-term resilience of these critical ecosystems.

## Data availability

All the climate data were sourced from WorldClim2 (http://www.worldclim.org/version2) using the GCS_WGS_1984 coordinate system at 30-arc sec (∼1 km2) resolutions, Resilient & Connected Networks (RCN) map from The Nature Conservancy ((https://maps.tnc.org/resilientland), the protected area information from the International Union for Conservation of Nature (IUCN) (http://www.cec.org/north-american-environmental-atlas/north-american-protected-areas-2021), and species occurrence points from the Global Biodiversity Information Facility (GBIF: www.gbif.org). These data will be made available on request.

## Acknowledgments

The authors thank numerous collaborators for their contribution to gathering the occurrence of grassland species. Special thanks to members of the Hamilton Lab for their help and support. Funding was provided by the Clean Water Land & Legacy Amendment in Minnesota, Doris Duke Charitable Foundation, Prairie Pothole Joint Venture, Wildlife Conservation Society’s Climate Adaptation Fund, The Nature Conservancy, and The Schatz Center for Tree Molecular Genetics at Pennsylvania State University.

## Authorship contribution statement

Santosh Kumar Rana: Writing – review & editing, Writing – original draft, Methodology, Formal analysis, Data curation, Conceptualization. Jessica Lindstrom: Writing – review & editing, Methodology, Data curation. Melissa A. Lehrer: Writing – review & editing. Marissa Ahlering: Writing – review & editing. Jill Hamilton: Writing – review & editing, Data curation, Conceptualization, Supervision.

