## Supplementary Figures for "Forecasting hotspots of grassland suitability under climate change for restoration"

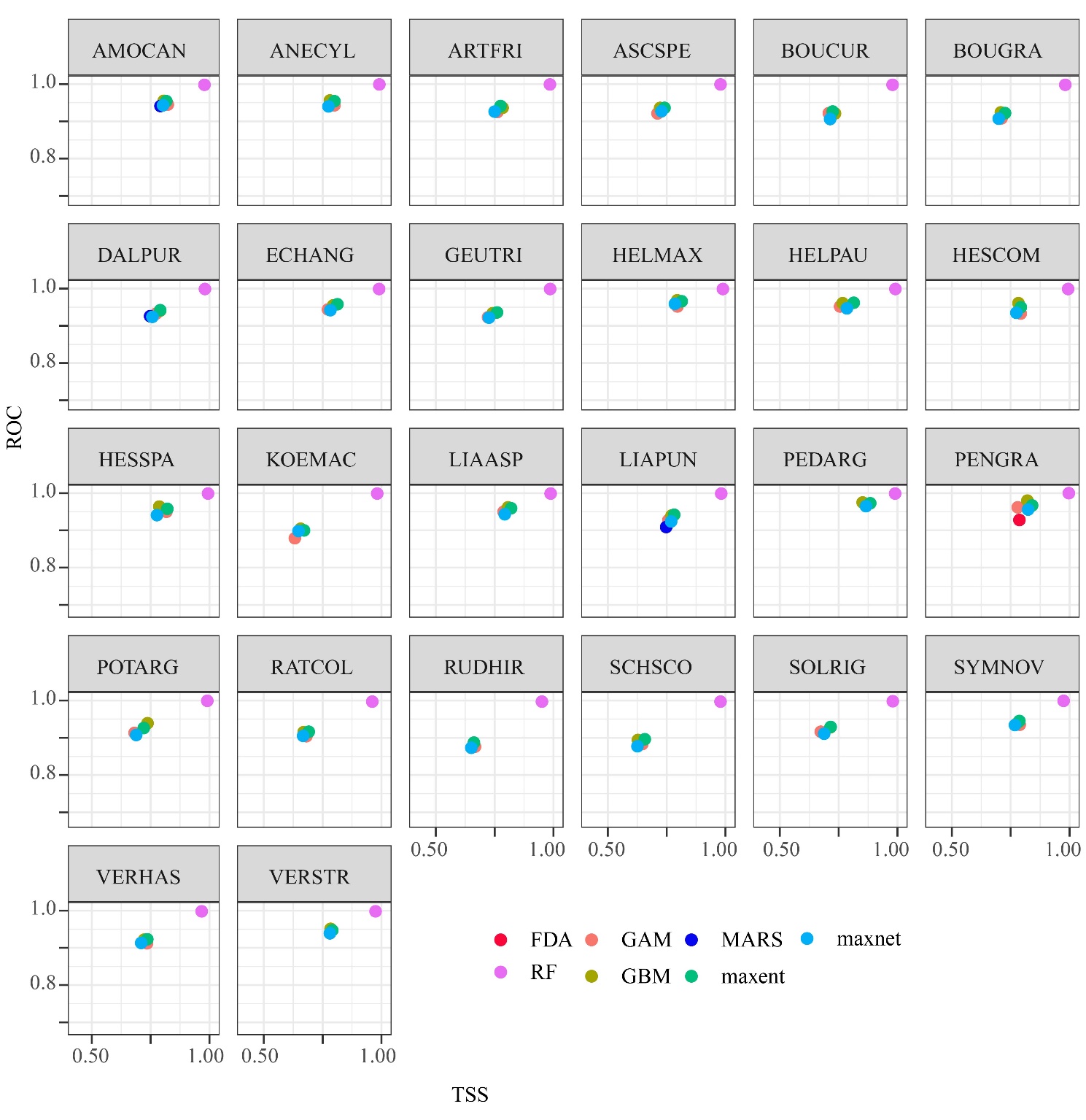
**Figure S1.** Performance statistics of all the calibrated algorithms in species distribution modeling using *Biomod2*. The plot of the mean (dot) of evaluation scores is used as the function of true skill statistics (TSS) (x-axis) versus area under the receiver operating characteristic curve (ROC) (y-axis). Refer Table S1 for the abbreviated species name.

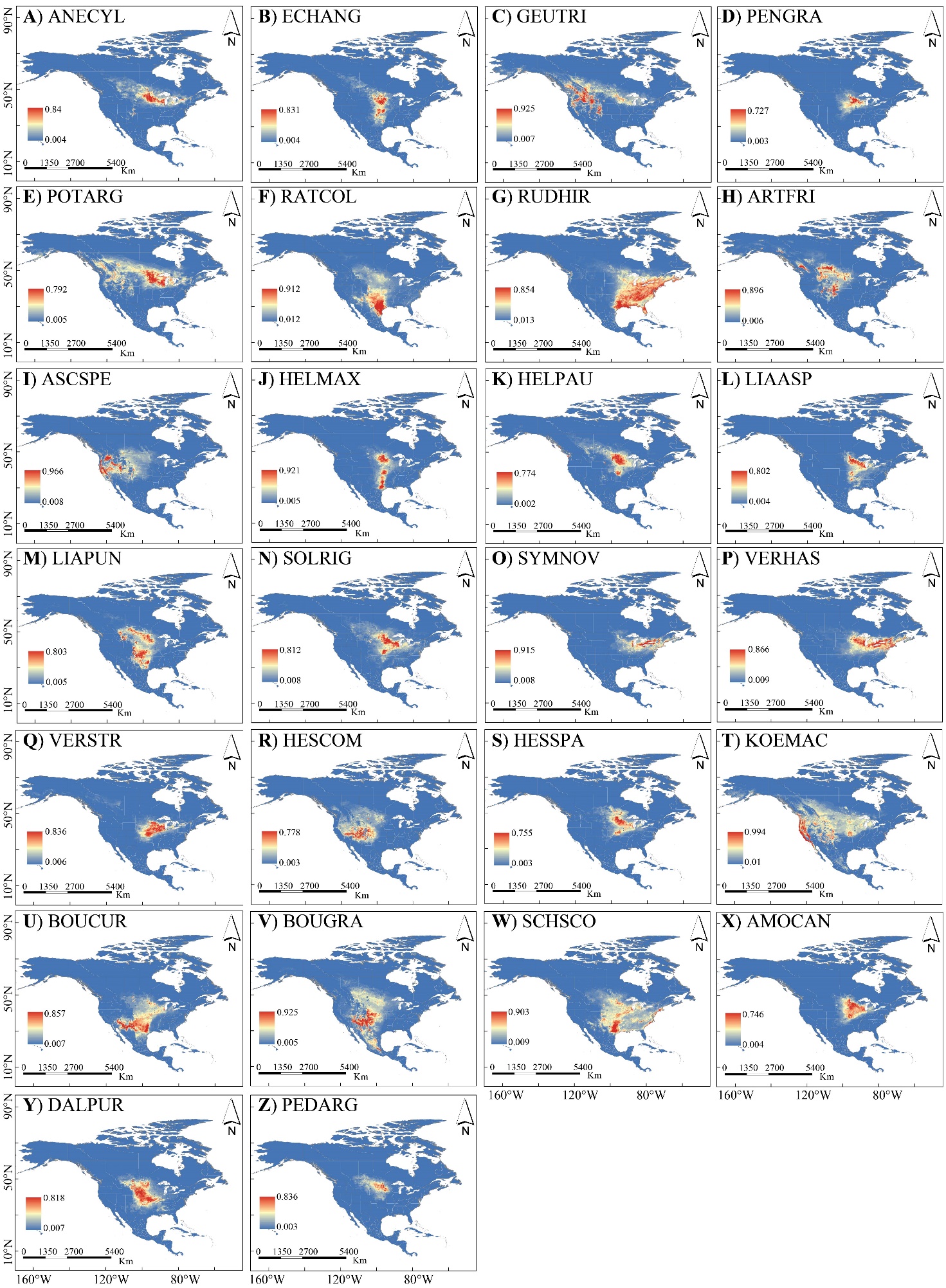
**Figure S2.** Grassland habitat suitability under the current bioclimatic variables scenario for 26 grassland species categorized as (**A**–**Q**) Forbs, (**R**–**W**) Grasses, and (**X**–**Z**) Legumes.

Refer Table S1 for the abbreviated species name.

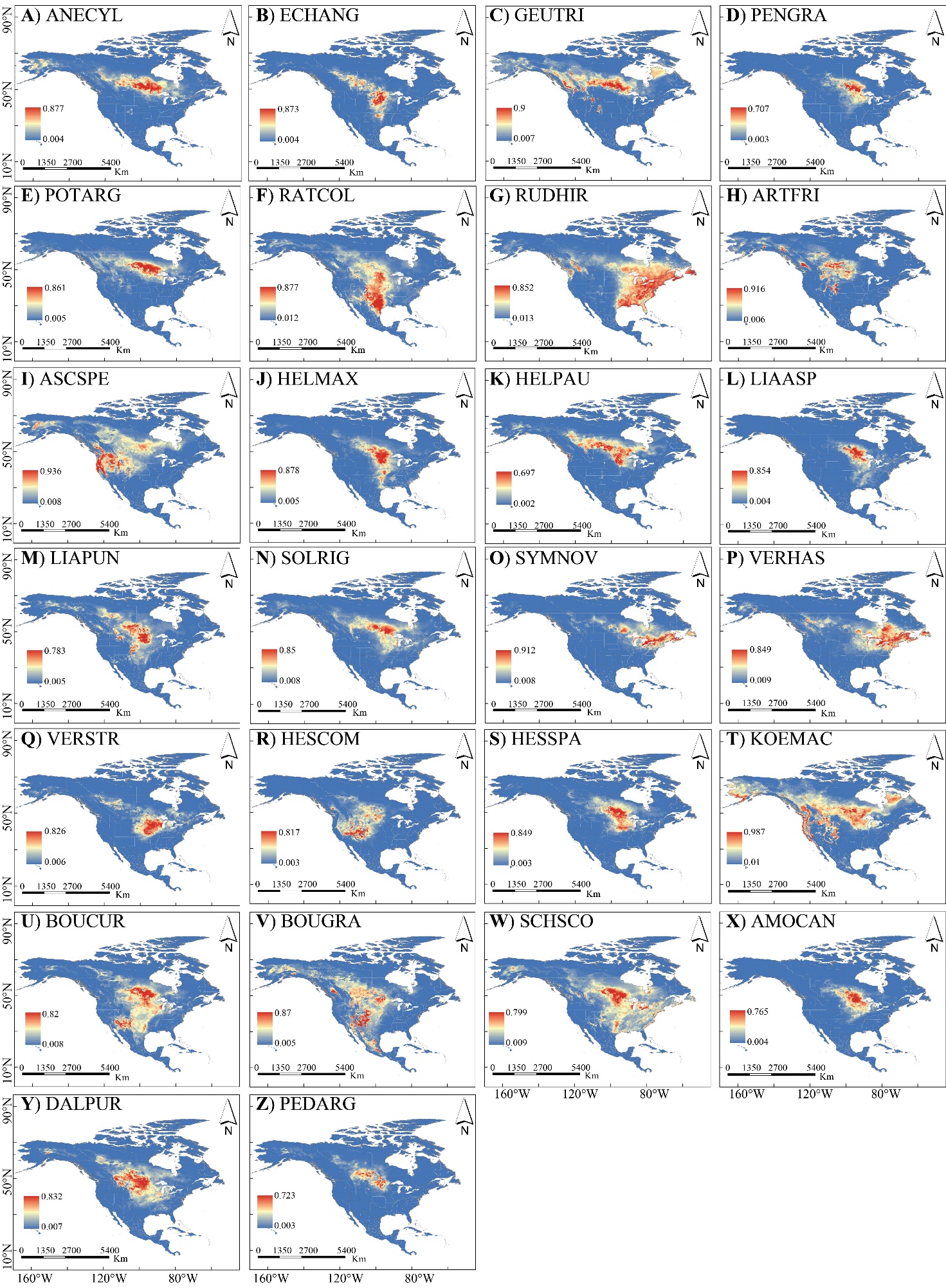
**Figure S3.** Grassland habitat suitability under the future bioclimatic variables scenario for 26 grassland species categorized as (**A**–**Q**) Forbs, (**R**–**W**) Grasses, and (**X**–**Z**) Legumes.

Refer Table S1 for the abbreviated species name.

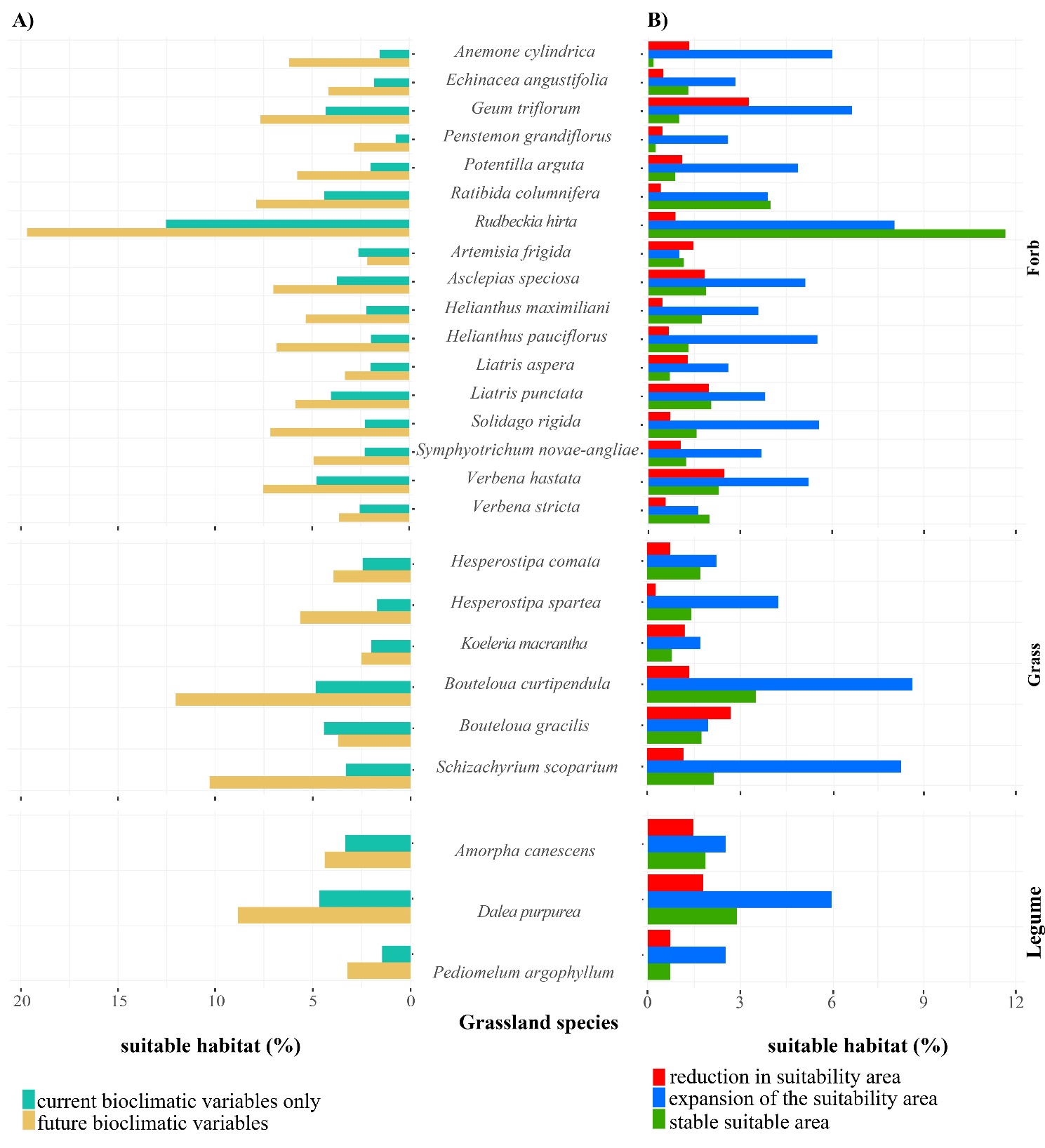

**Figure** **S4.** Predicted suitable areas of functionally categorized 26 grassland species (Forbs, Grasses, and Legumes) under **A**) current and future bioclimatic variables’ scenarios, and **B**) the categorized reduced, expanded, and stable areas calculated as a function of raster aggregation for the current and future bioclimatic variables’ eSDM projections. ‘Reduction’ represents a decrease, ‘Expansion’ represents an increase, and ‘Stable’ represents no change in delineated suitable habitat area in the future (2050) as compared to the current scenario projected through eSDM of bioclimatic variables. The bar graph represents the percentage (%) of suitable habitat area out of the total modeled area (total area of North America i.e., 24.71 million km²).

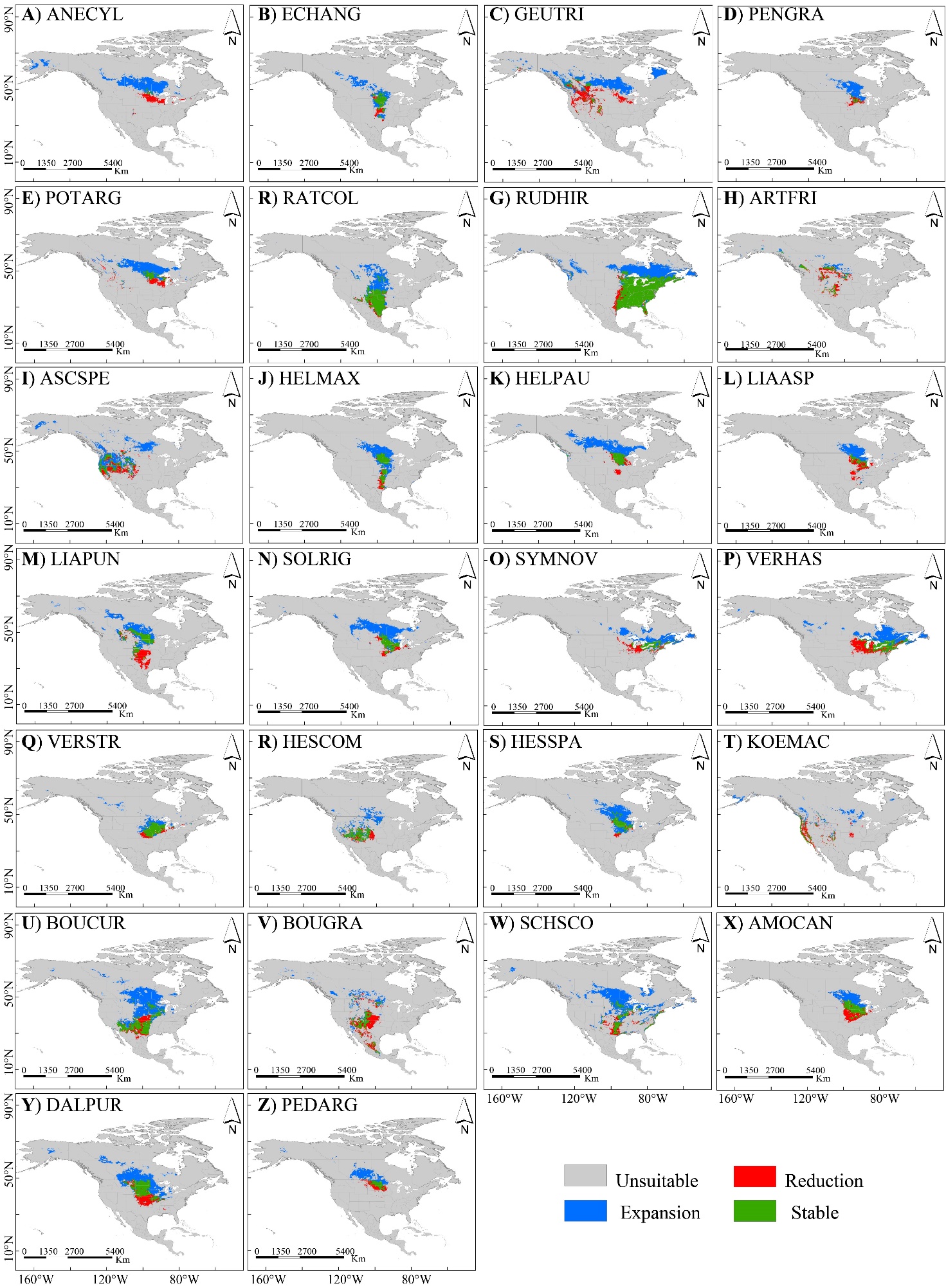
**Figure S5.** Grassland habitat suitability classified as the reduced, stable, and expanded areas for 26 grassland species categorized as (**A**–**Q**) Forbs, (**R**–**W**) Grasses, and (**X**–**Z**) Legumes. Refer Table S1 for the abbreviated species name.
